# Gaze and Movement Assessment (GaMA): Inter-site validation of a visuomotor upper limb functional protocol

**DOI:** 10.1101/681437

**Authors:** Heather E. Williams, Craig S. Chapman, Patrick M. Pilarski, Albert H. Vette, Jacqueline S. Hebert

## Abstract

**Background:** Successful hand-object interactions require precise hand-eye coordination with continual movement adjustments. Quantitative measurement of this visuomotor behaviour could provide valuable insight into upper limb impairments. The Gaze and Movement Assessment (GaMA) was developed to provide protocols for simultaneous motion capture and eye tracking during the administration of two functional tasks, along with data analysis methods to generate standard measures of visuomotor behaviour. The objective of this study was to investigate the reproducibility of the GaMA protocol across two independent groups of non-disabled participants, with different raters using different motion capture and eye tracking technology.

**Methods:** Twenty non-disabled adults performed the Pasta Box Task and the Cup Transfer Task. Upper body and eye movements were recorded using motion capture and eye tracking, respectively. Measures of hand movement, angular joint kinematics, and eye gaze were compared to those from a different sample of twenty non-disabled adults who had previously performed the same protocol with different technology, rater and site.

**Results:** Participants took longer to perform the tasks versus those from the earlier study, although the relative time of each movement phase was similar. Measures that were dissimilar between the groups included hand distances travelled, hand trajectories, number of movement units, eye latencies, and peak angular velocities. Similarities included all hand velocity and grip aperture measures, eye fixations, and most peak joint angle and range of motion measures.

**Discussion:** The reproducibility of GaMA was confirmed by this study, despite a few differences introduced by learning effects, task demonstration variation, and limitations of the kinematic model. The findings from this study provide confidence in the reliability of normative results obtained by GaMA, indicating it accurately quantifies the typical behaviours of a non-disabled population. This work advances the consideration for use of GaMA in populations with upper limb sensorimotor impairment.

## Introduction

Various sensorimotor impairments including stroke [1], amputation [2], and spinal cord injury [3] lead to deficits in upper limb performance that can hamper activities of daily living requiring precise hand-object interactions [4]. Various functional assessments are used to gauge the functional impact of upper limb impairment and to monitor rehabilitative progress thereafter [5], [6]. However, such assessments often do not precisely quantify hand and joint movements, grip adjustments [7], [8], or hand-eye interaction, which is recognized as an important behaviour during grasp control [9], [10]. Quantitative measurement of visuomotor behaviour collected during the execution of functional tasks can enhance the understanding of these movement features. Measurement technologies commonly used for this purpose include eye tracking and motion capture. Assessments reliant on such specialized equipment, however, suffer from a lack of standardized protocols and can be criticized as not being generalizable to activities of daily function. Furthermore, technology-based assessments risk becoming obsolete as newer technologies emerge, hindering the opportunity for robust comparisons over time.

The Gaze and Movement Assessment (GaMA) protocol was designed to overcome these limitations. GaMA includes two standardized functional upper limb tasks that incorporate common dextrous hand demands of daily living [7]. GaMA also includes an analysis software, which requires a standardized data set of synchronized motion and eye data coordinates as input (obtained using motion capture and eye tracking during functional task execution), and outputs metrics of hand movement, angular joint kinematics, and eye gaze behavior [7]–[9]. GaMA’s input data set can be obtained by various data collection hardware and software solutions, rendering the assessment protocol amenable to technological evolution (for example, markerless motion capture and mobile eye trackers). Additionally, GaMA measures remain relevant and equipment-independent for future comparative purposes, potentially both within and across research sites. The ability to compare results across sites would be extremely valuable as it could facilitate larger subgroup comparisons when smaller populations of individuals with upper limb impairments are studied, such as upper limb prosthesis users.

In order to validate a new protocol such as GaMA, it is essential to determine reproducibility. Reproducibility of a test or method is defined as the closeness of the agreement between independent results obtained by following the same procedures, but under different experimental conditions [11]. Due to the inherent variability found in clinical populations, reproducibility of a test to assess movement behaviour is typically first studied in a non-disabled population. While intra-rater test-retest reliability of GaMA has been demonstrated for hand movement and angular joint kinematic measures for non-disabled individuals [7], [8], it has yet to be determined whether these and other measures obtainable by GaMA are reproducible across raters and sites. Furthermore, it is often assumed that the non-disabled population will behave similarly (or identically) across test sites; yet, it is known that deviations from protocols can result in data set disparity amongst the population [12]. If a standardized protocol can be shown to yield measures that are similar across sites, the data sets could be combined for a richer understanding (or more saturated data set) of non-disabled movement behaviour.

The objective of this study, therefore, was to conduct an inter-site validation of GaMA by assessing the reproducibility of the visuomotor measures in non-disabled individuals presented by Valevicius et al. and Lavoie et al. [7]–[9]. More specifically, this study sought to determine whether the same hand movement, angular joint kinematic, and eye gaze measures could be obtained by testing a second independent group of non-disabled participants, at a different site equipped with different motion capture and eye tracking technology, and administered by a different rater. Establishing the reproducibility of GaMA in the non-disabled population advances its consideration as an outcome assessment protocol for populations with sensorimotor impairments of the upper limb.

## Methods

For comparative purposes, the research conducted by Valevicius et al. [7], [8] and Lavoie et al. [9] is referred to in this paper as ‘the original study’, and the data set analyzed by these studies is referred to as ‘the original data set’. The new research presented in this article is referred to as ‘the repeated study’ and its data as ‘the repeated data set’. Unless otherwise specified, the same procedures were followed in both studies. Ethical approval for these procedures was obtained by the University of Alberta Health Research Ethics Board (Pro00054011), the Department of the Navy Human Research Protection Program, and the SSC-Pacific Human Research Protection Office.

### Participants

A total of 22 non-disabled adults were recruited to participate in the repeated study. Data from two participants were removed due to problems arising from software issues. The characteristics of the 20 participants from the original study [7]–[9] and the 20 participants in the repeated study are detailed in **Error! Reference source not found.**. In both studies, two participants performed the tasks without corrected vision, since they had to remove their glasses to don the eye tracker. These participants, however, reported that their vision was sufficient to allow them to confidently perform the task.

**Table 1:** Original and repeated study participant characteristics.

| Research Participant Characteristics | Original Study | Repeated Study |
| --- | --- | --- |
| Male participants | 11 | 13 |
| Female participants | 9 | 7 |
| Self-reported right-handed participants | 18 | 19 |
| Participants with normal or corrected to normal vision | 18 | 18 |
| Participant age (years – mean ± standard deviation) | 25.8 ± 7.2 | 24.4 ± 7.3 |
| Participant height (cm – mean ± standard deviation) | 173.8 ± 8.3 | 171.0 ± 7.7 |

### Equipment

Motion capture and eye tracking hardware and software specifications for the original study and the repeated study are indicated in **Error! Reference source not found.**. The equipment was set up in the repeated study as specified in the original study [7]–[9]. Rigid plates and a headband (each holding four retroreflective markers) were attached to the participant in accordance with Boser et al.’s *Clusters Only* kinematic model [13]. To improve rigid body motion tracking in the repeated study, the hand plates were redesigned as shown in Fig 1. For both studies, markers were attached to the index finger (middle phalange) and thumb (distal phalange) [7]; a head-mounted eye tracker was placed on the participant and positioned in accordance with the manufacturer’s instructions; and a motion capture calibration pose was collected for each participant, as outlined by Boser et al. [13].

**Table 2:** Specifications of the motion capture and eye tracking systems used in the original and repeated studies.

| Specifications | Original Study | Repeated Study |
| --- | --- | --- |
| Motion capture camera | Vicon Bonita 10<br>(Vicon Motion Systems Ltd,<br>Oxford, UK) | OptiTrack Flex 13<br>(Natural Point, OR, USA) |
| Number of cameras | 12 | 8 |
| Camera sampling frequency | 120 Hz | 120 Hz |
| Head-mounted binocular eye tracker | Dikablis Professional 2 (Ergoneers GmbH, Manching, Germany) | Pupil (Pupil Labs GmbH, Berlin, Germany) |
| Eye camera sampling frequency | 60 Hz | 120 Hz |

**Fig 1.**
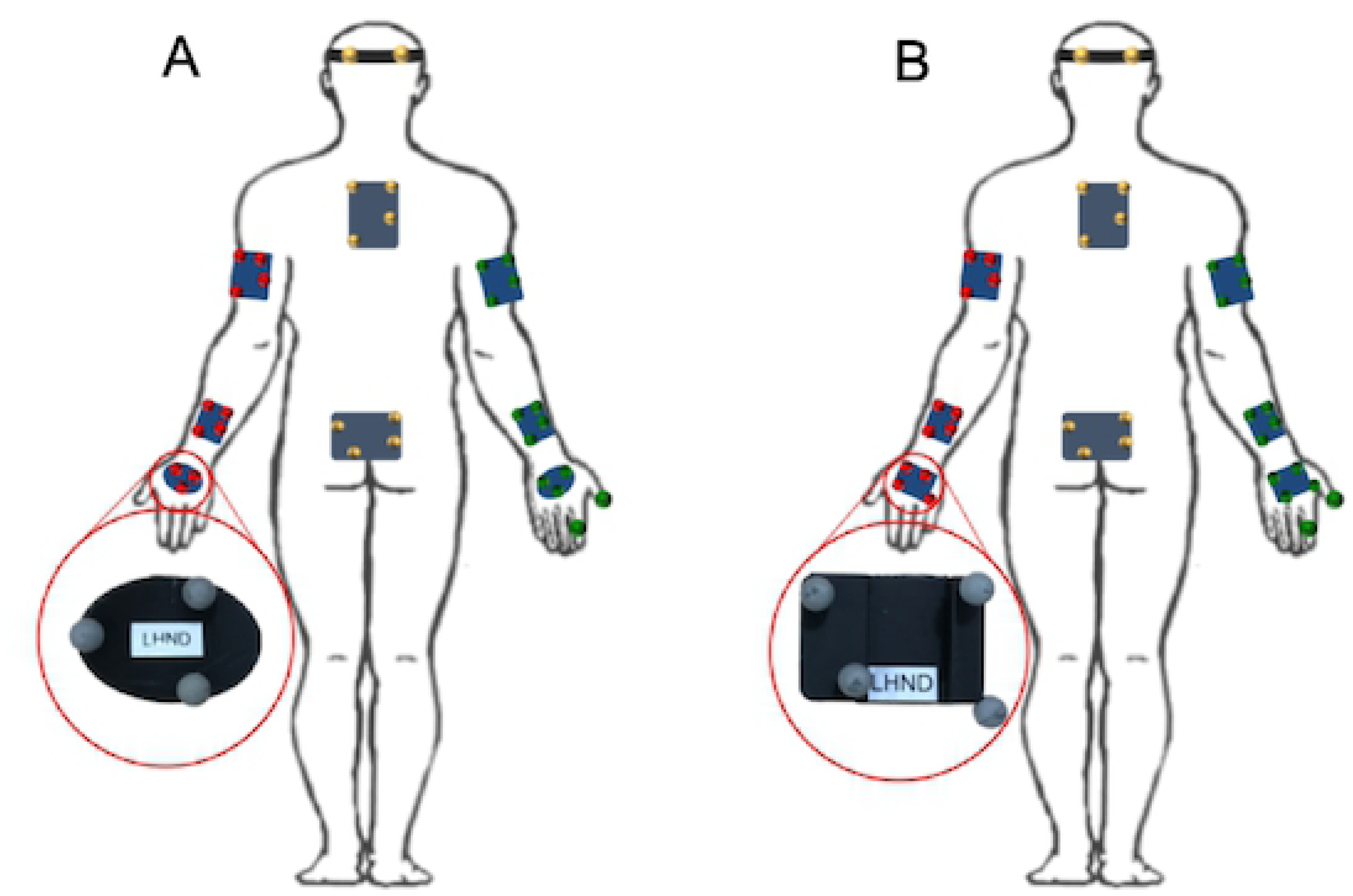
Retroreflective marker placement. Marker placement for participants in the original study (A) and repeated study (B), showing differences in the hand marker plate designs.

### Data Collection

In both studies, the two functional tasks introduced by Valevicius et al. (the Pasta Box Task and Cup Transfer Task) [7] were administered. Each participant completed 20 error-free trials of the two tasks, while simultaneous motion and eye tracking data were collected. Prior to this, each participant was given verbal instructions, a demonstration, and at least one familiarization trial of each functional task. Task order was randomized for each participant. At least two gaze calibrations (outlined by Lavoie et al. [9]) were collected before participants executed their initial trial of each task, and one after they completed their final trial of the last task; given that there were two functional tasks, a minimum of 5 calibrations were performed per participant.

The original data collection protocol differed from the repeated study in one notable way. In the original study, every participant performed a total of 60 trials of each task, 20 of which were under each of the following conditions: (1) only motion capture data were collected, (2) only eye tracking data were collected, and (3) both motion capture and eye tracking data were collected. As the repeated study consisted solely of collecting data during simultaneous motion capture and eye tracking, it was only compared to that of the original data set captured under condition (3) ‘both’. In the original study, the order of conditions for each participant was block randomized to one of 4 block orders, with motion (1) and both (3) conditions always sequential. As a consequence of the partial randomization order, three quarters of the original study participants were afforded at least 20 extra trials executing each functional task prior to testing under the ‘both’ condition.

### Experimental Data Analysis

Data analysis in the repeated study was undertaken as outlined by Valevicius et al. and Lavoie et al. [7]–[9]: motion capture marker trajectory data and pupil position data were filtered and synchronized; hand movement and angular kinematic measures were calculated; the virtual location of the participant’s gaze (represented by a gaze vector) was determined using gaze calibration data; and gaze fixations to areas of interest were calculated. Due to insufficient pupil data, the data from one participant were removed from the repeated data set for the Cup Transfer Task, and data from four participants were removed for the Pasta Box Task.

For each functional task, the repeated data set were divided into distinct *movements* based on hand velocity, the velocity of the task object(s), and grip aperture values, as per Valevicius et al. [7]. The data from each movement were further segmented into the *phases* of ‘Reach’, ‘Grasp’, ‘Transport’, ‘Release’, and ‘Home’; the Home phase was not used for data analysis. Due to the short duration of the Grasp and Release phases, combined *movement segments* of ‘Reach-Grasp’ and ‘Transport-Release’ were used in hand movement analysis [7]. Eye latency measures were calculated at instances of *phase transition*, both at the end of a Grasp phase and at the beginning of a Release phase (referred to as ‘Pick-up’ and ‘Drop-off’ by Lavoie et al. [9]). An illustration of how one distinct movement was separated into the abovementioned subsets (phases, movement segments, and phase transitions) can be found in Fig 2.

**Fig 2.**
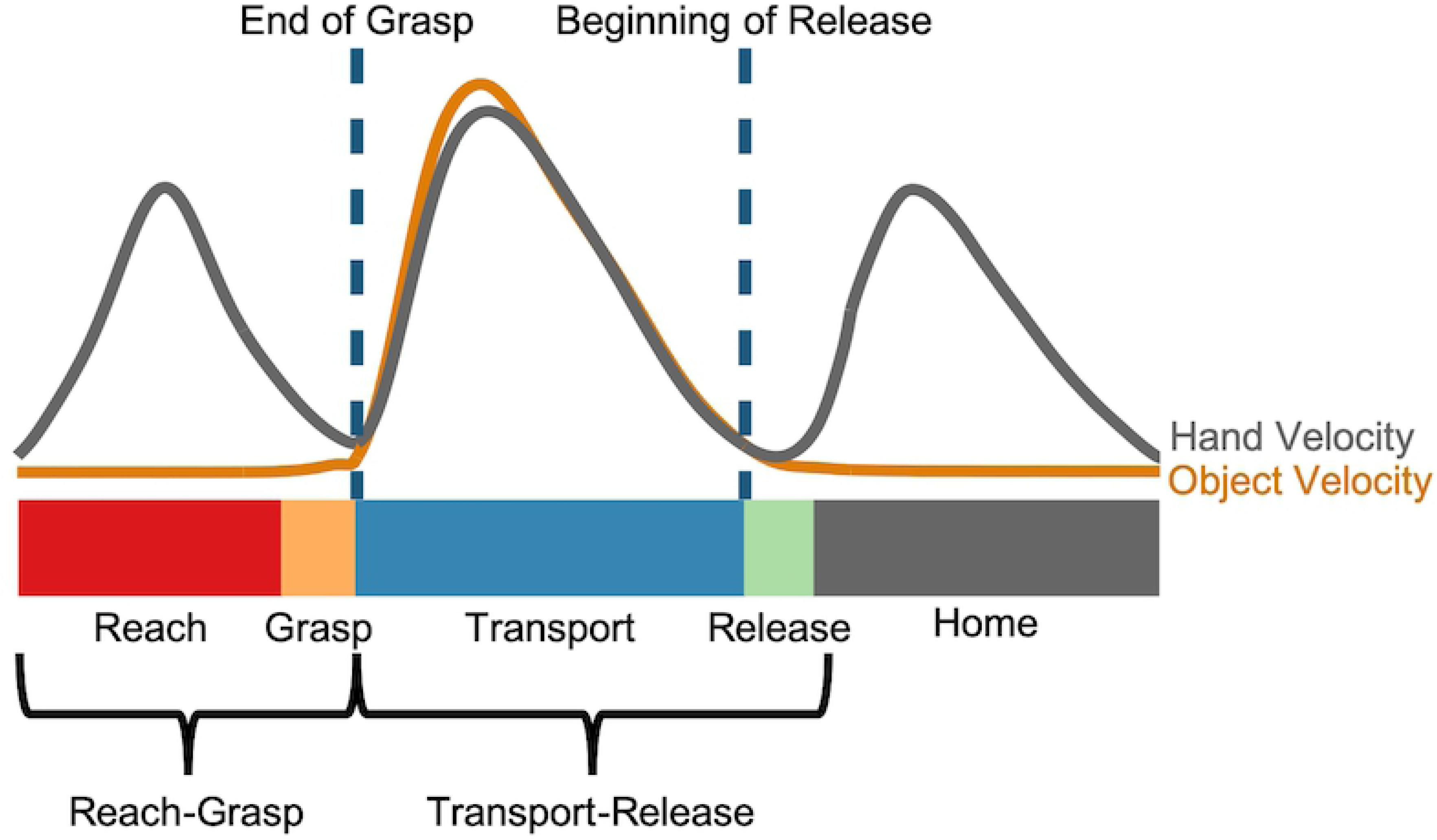
Phase transitions, phases, and movement segments within one movement. Typical hand and object velocity profiles are displayed in grey and orange lines, respectively. Reach, Grasp, Transport and Release phases are presented along the bar as red, orange, blue and green, respectively. Home (grey bar) refers to the standardized location to which the hand returns at the completion of the movement.

### GaMA Measures

Duration (phase and trial), hand movement, angular joint kinematic, and eye gaze measures were calculated for the original and repeated studies, as outlined by Valevicius et al. [7], [8] and Lavoie et al. [9], and are listed in **Error! Reference source not found.**. Lavoie et al.’s ‘fixations to future’ measure was not considered in this study as these fixations were shown to be unlikely to occur in non-disabled participants for both tasks [9]. In addition to the measures listed in **Error! Reference source not found.**, the relative duration of each phase was calculated as the percent of time spent in that phase, relative to the given Reach-Grasp-Transport-Release sequence.

**Table 3:** Comparative measures, including duration, hand movement, angular joint kinematic, and eye gaze measures, and the subsets of each movement for which they were calculated.

| Type of Measure | Measures | Movement Subsets |
| --- | --- | --- |
| Duration (from Lavoie et al. [9]) | Phase duration | Reach, Grasp, Transport, Release |
| Hand movement (from Valevicius et al. [7]) | Hand distance travelled<br>Hand trajectory variability<br>Peak hand velocity<br>Percent-to-peak hand velocity<br>Number of movement units | Reach-Grasp, Transport-Release |
|  | Peak grip aperture<br>Percent-to-peak grip aperture<br>Percent-to-peak hand deceleration | Reach-Grasp |
|  | Percent fixation to Hand in Flight<br>Number of fixations to Hand in Flight | Reach, Transport |
|  | Eye Arrival Latency<br>Eye Leaving Latency | End of Grasp, Beginning of Release |
| Angular joint kinematics (from Valevicius et al. [8]) | Peak angle, range of motion, and peak angular velocity for the following degrees of freedom:<br>• Trunk flexion/extension<br>• Trunk lateral bending<br>• Trunk axial rotation<br>• Shoulder flexion/extension<br>• Shoulder abduction/adduction<br>• Shoulder internal/external rotation<br>• Elbow flexion/extension<br>• Forearm pronation/supination<br>• Wrist flexion/extension<br>• Wrist ulnar/radial deviation | Movement only |
| Eye gaze (from Lavoie et al. [9]) | Percent fixation to Current<br>Number of fixations to Current | Reach, Grasp, Transport, Release |

In the repeated study, the calculation of hand movement measures was altered due to the creation of a virtual rectangular prism, which approximated the participant’s hand position at each point in time. Using the centre of this prism, hand position and velocity were subsequently calculated. For comparative purposes, the original study’s hand movement results were recalculated via this methodology rather than the original calculation of Valevicius et al. using the average position of the three hand plate markers [7]).

### Statistical Analysis

The aim of the statistical analysis was to detect significant differences between the original and repeated data sets, and to determine whether such differences were more pronounced for particular movements and/or movement subsets (phase, movement segment, or phase transition). To investigate differences between the two groups of participants, a series of repeated-measures analyses of variance (RMANOVAs) and pairwise comparisons were conducted for each measure and task. RMANOVA group effects or interactions involving group were followed up with either an additional RMANOVA or pairwise comparisons between groups if the Greenhouse-Geisser corrected *p* value was less than 0.05. Pairwise comparisons were considered to be significant if the Bonferroni corrected *p* value was less than 0.05. Detailed statistical analysis methods can be found in supplementary materials (S1 Text).

## Results

### Duration

For both the Pasta Box Task (or ‘Pasta’) and the Cup Transfer Task (or ‘Cups’), the repeated study participants took significantly more time to complete the tasks than the original study participants (Pasta: 11.8 ± 3.4 seconds versus 8.8 ± 1.2 seconds, p < 0.01; Cups: 13.9 ± 2.5 seconds versus 10.5 ± 1.3 seconds, p < 0.0001). The repeated study participants had longer phase durations than the original study participants, with all Grasp and Transport phases and the Movement 2 Release phase significantly prolonged in Pasta, and all phases significantly prolonged in Cups (S2 Table). The two participant groups, however, displayed similar relative phase durations throughout both tasks, with no significant differences.

### Hand Movement

The repeated study participants had greater hand distances travelled than the original study participants, with significant increases in Movement 1 & 3 segments of Pasta (S3 Table) and in all Cups movement segments, except for Movement 1 & 4 Transport-Releases (S4 Table). However, Fig 3 (Pasta) and Fig 4 (Cups) show that the average hand trajectories chosen by both participant groups were similar. The repeated study participants also had larger hand trajectory variability than the original study participants, with significant increases in all Pasta movement segments except for Movement 3 Transport-Release (S4 Table) and all Cups movement segments (S5 Table). The repeated study participants had a greater number of movement units than the original study participants, with significant increases in all movement segments of Pasta and for Movement 1 & 4 Reach-Grasp and Movement 1 to 3 Transport-Release segments of Cups.

**Fig 3.**
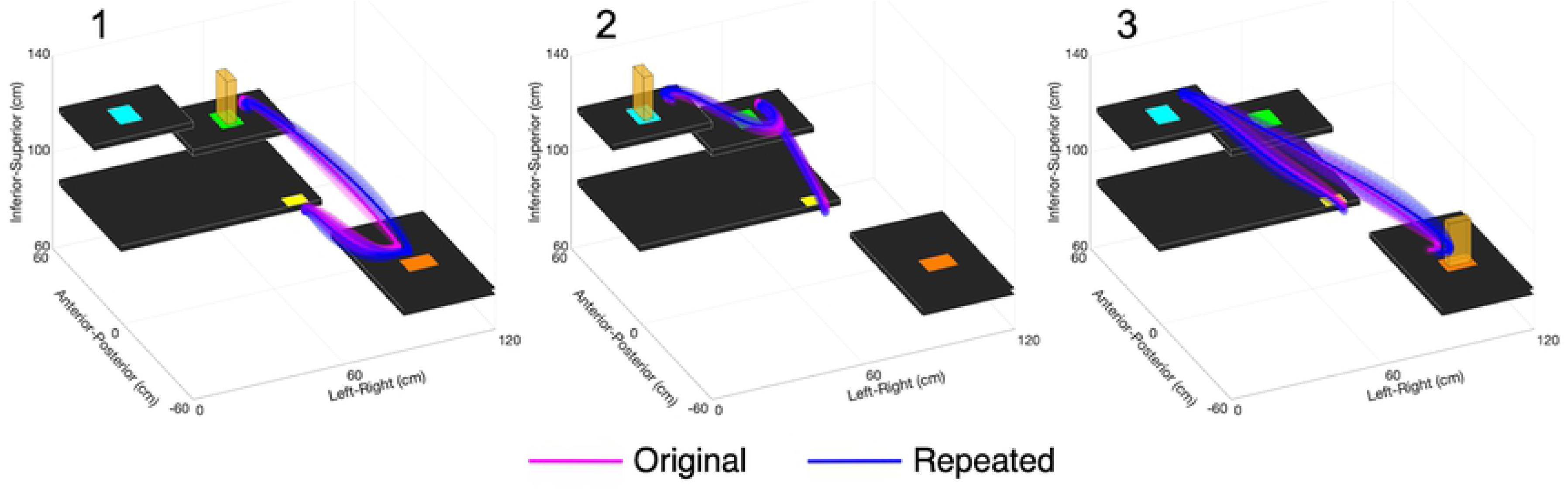
Pasta Box Task hand trajectories. Trajectories are displayed for participants in the original (pink) and repeated (blue) studies for Movements 1, 2, and 3. The solid lines represent participant group averages, and the three-dimensional shading represents the standard deviation of participant group means.

**Fig 4.**
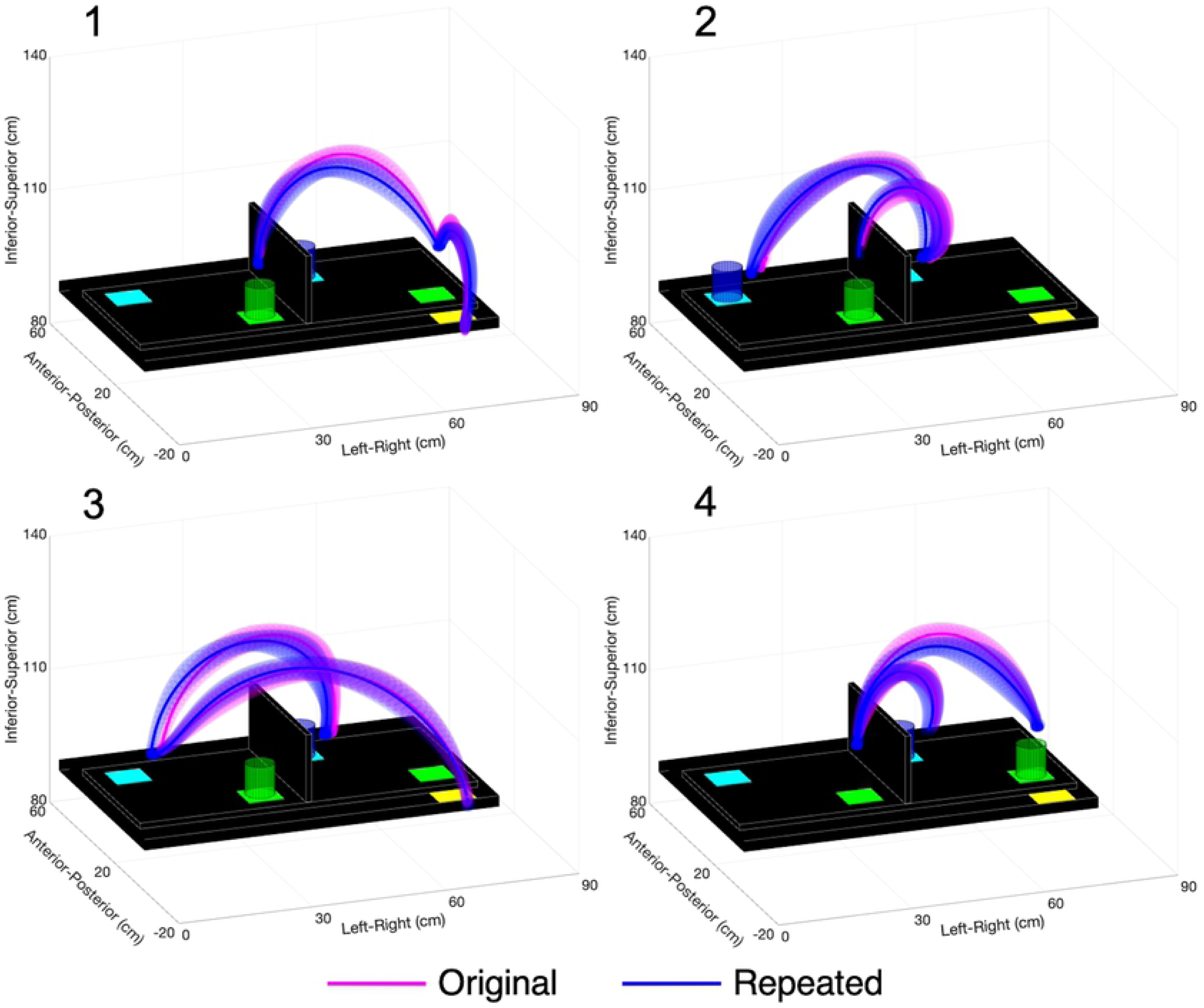
Cup Transfer Task hand trajectories. Trajectories are displayed for participants in the original (pink) and repeated (blue) studies for Movements 1, 2, 3, and 4. The solid lines represent participant group averages, and the three-dimensional shading represents the standard deviation of participant group means.

Participants in the original and repeated studies had similar hand velocity profiles for both tasks, as shown in Fig 5A and 5B. Although the peaks in the repeated study appeared smaller, these differences were non-significant throughout both tasks (S4 Table and S5 Table). Significant percent-to-peak hand velocity differences were identified for the Movement 1 Reach-Grasp segment of Pasta and the Movement 2 & 3 Reach-Grasp segments of Cups, but the differences between the mean values of the two participant groups were less than one standard deviation of the original study results. Participants in the original and repeated studies showed similar percent-to-peak hand deceleration values, with no significant differences in Pasta and a significantly difference only for the Movement 4 Reach-Grasp segment of Cups. However, the difference between the mean values of the two participant groups in this movement segment was less than one original study standard deviation.

**Fig 5.**
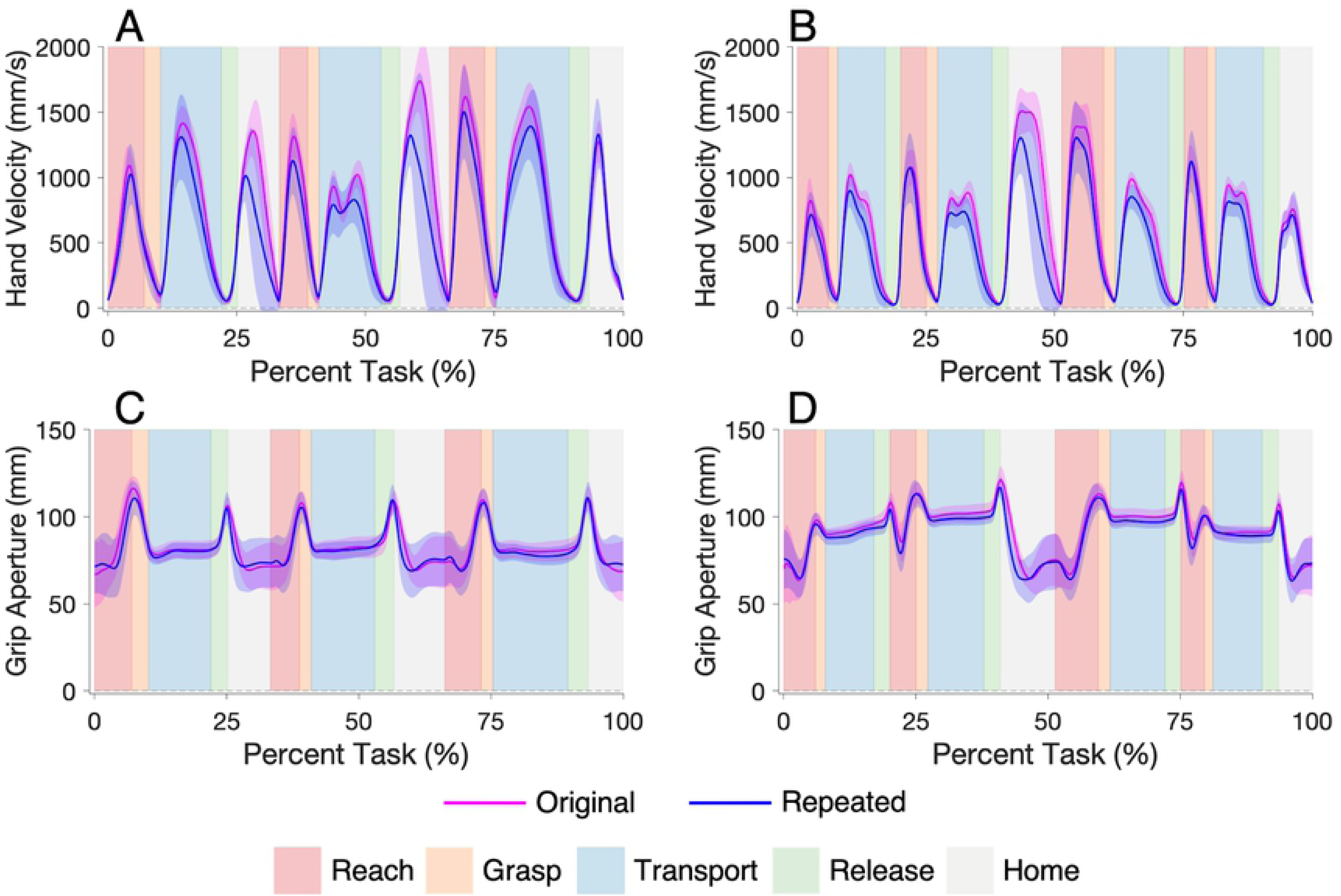
Hand velocity profiles for the Pasta Box Task (A) and the Cup Transfer Task (B); and grip aperture profiles for the Pasta Box Task (C) and the Cup Transfer Task (D). Original study data are presented in pink, and repeated study data in blue. The solid lines represent participant group averages, and the shading represents one standard deviation of the participant group means. This task is segmented into Reach (red), Grasp (orange), Transport (blue), Release (green), and Home (grey) phases for each movement.

Participants in the original and repeated studies had similar grip aperture profiles for both tasks, as shown in Fig 5C and 5D, with no significant differences in peak grip aperture identified for either task. Also, no significant differences in percent-to-peak grip aperture were identified in Pasta, and a significant difference was only identified in the Movement 4 Reach-Grasp segment of Cups.

### Angular Joint Kinematics

Angular kinematic trajectories illustrating the average joint trajectories of participants are shown in Fig 6 (Pasta) and Fig 7 (Cups). Similar angular kinematic profiles existed between the original and repeated study participants, with only a few differences; participants in the repeated study had an increased standard deviation for trunk flexion/extension (both tasks), and an offset was present between the wrist flexion/extension angles (both tasks) and between the wrist ulnar/radial deviations angles (Pasta only) of the two participant groups. Angular kinematic measures are presented in **Error! Reference source not found.** (Pasta) and **Error! Reference source not found.** (Cups). The original and repeated study participants generally had similar peak joint angles in both tasks. Significant peak angle differences were found in wrist flexion/extension for Movements 1 and 2 of Pasta and all movements of Cups, and in wrist radial/ulnar deviation for all movements of Pasta.

**Fig 6.**
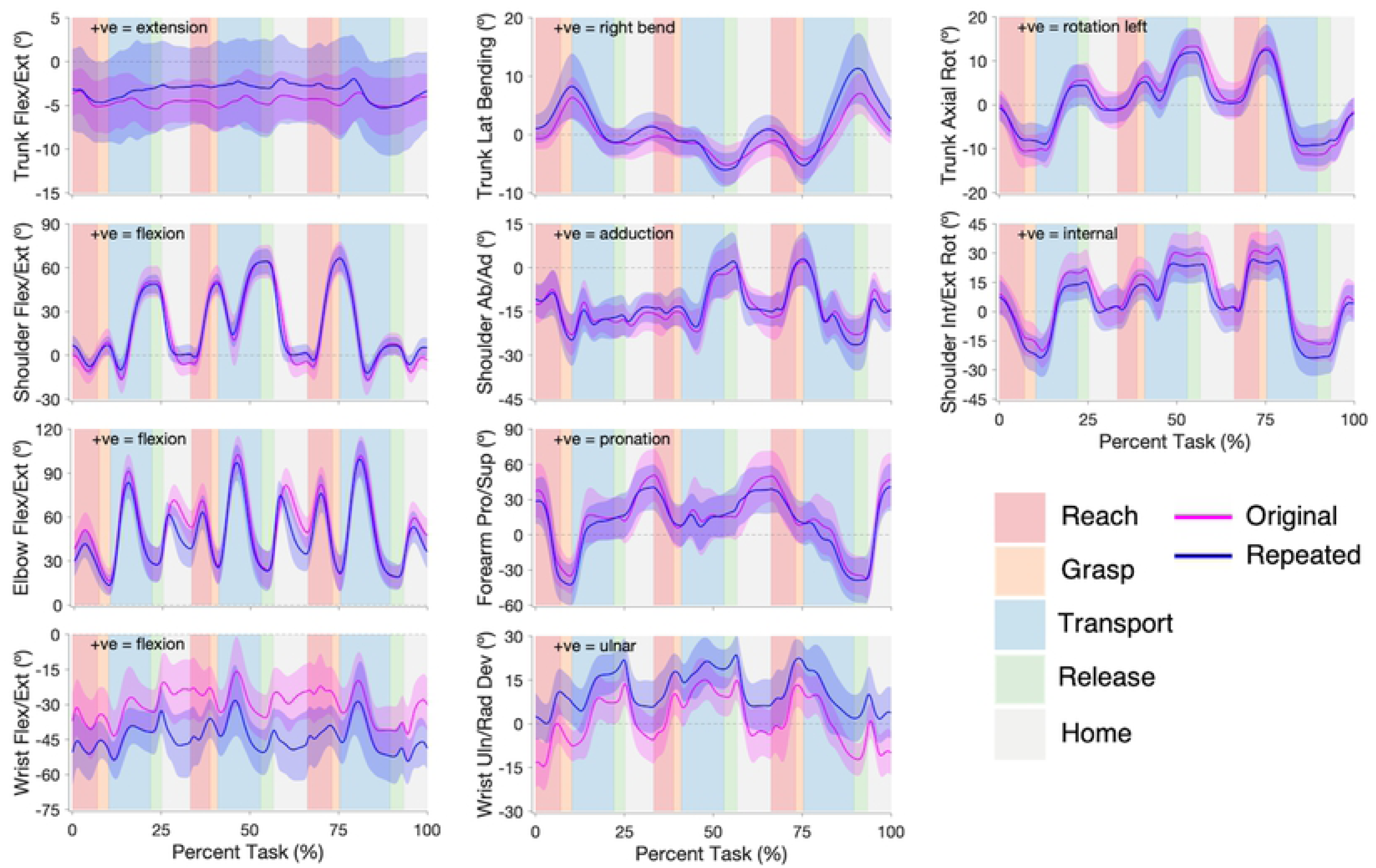
Pasta Box Task angular joint trajectories. Original (pink) and repeated (blue) studies angular joint trajectories for trunk flexion/extension, lateral bending, and axial rotation; shoulder flexion/extension, abduction/ adduction, and internal/external rotation; elbow flexion/extension and forearm pronation/supination; and wrist flexion/extension and ulnar/radial deviation. The solid lines represent participant group averages, and the shading represents one standard deviation of the participant group means. The task is segmented into Reach (red), Grasp (orange), Transport (blue), Release (green), and Home (grey) phases for each movement.

**Fig 7.**
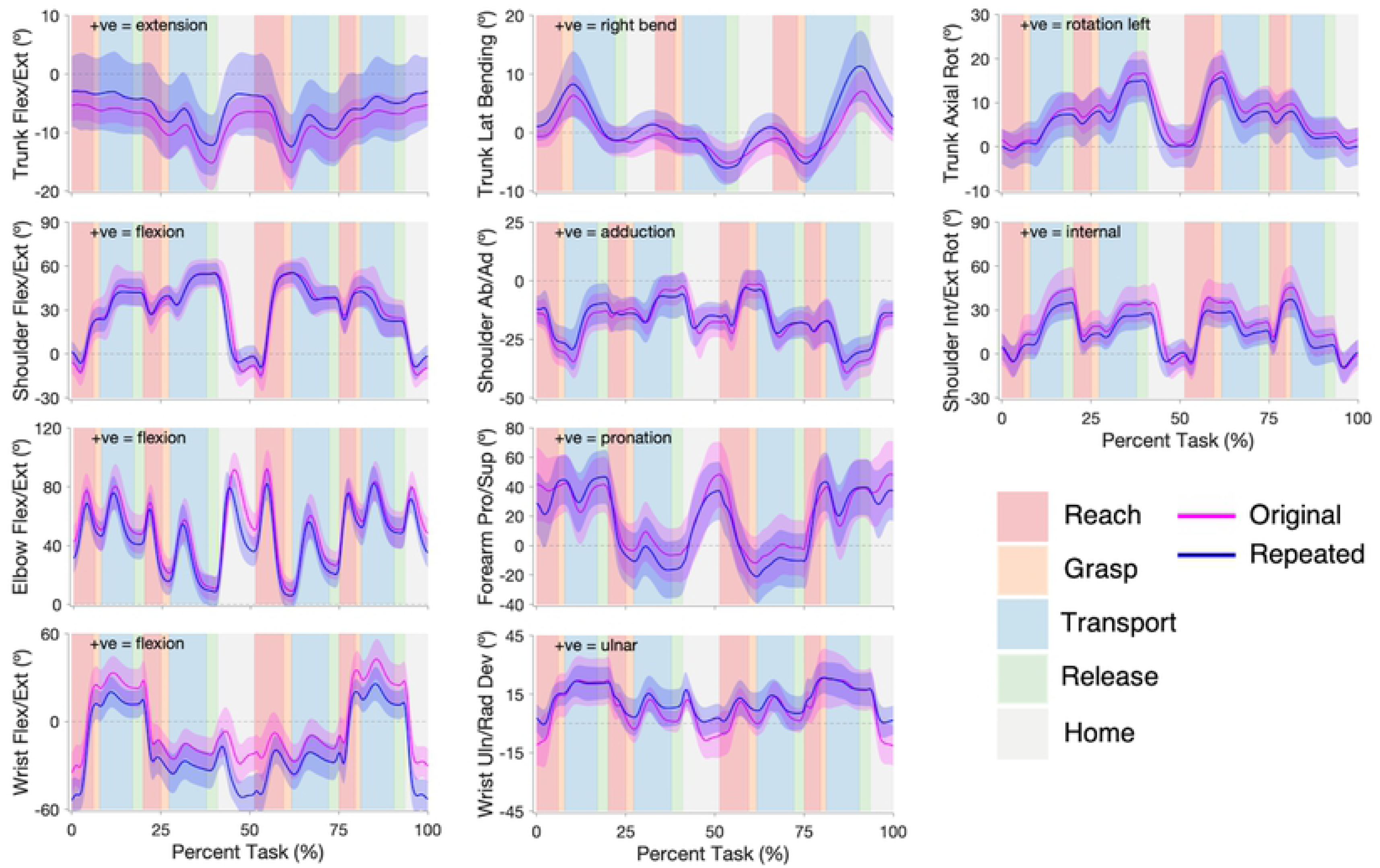
Cup Transfer Task angular joint trajectories. Original (pink) and repeated (blue) studies angular joint trajectories for trunk flexion/extension, lateral bending, and axial rotation; shoulder flexion/extension, abduction/ adduction, and internal/external rotation; elbow flexion/extension and forearm pronation/supination; and wrist flexion/extension and ulnar/radial deviation. The solid lines represent participant group averages, and the shading represents one standard deviation of the participant group means. The task is segmented into Reach (red), Grasp (orange), Transport (blue), Release (green), and Home (grey) phases for each movement.

**Table 4:** Pasta Box Task angular joint kinematic values, with significant results of the RMANOVAs and pairwise comparisons.

|  | M | Peak Angle (degrees) |  |  | Range of Motion (degrees) |  |  | Peak Angular Velocity (degrees/s) |  |  |
| --- | --- | --- | --- | --- | --- | --- | --- | --- | --- | --- |
|  |  | *p* | Original | Repeated | *p* | Original | Repeated | *p* | Original | Repeated |
| Trunk FE | 1 | ns | -2.1 ± 2.4 | -0.9 ± 4.7 | ns | 4.9 ± 1.6 | 6.0 ± 2.2 | ns | 18.8 ± 5.4 | 23.7 ± 7.8 |
|  | 2 | ns | -2.7 ± 2.6 | 0.2 ± 5.0 | * | 3.6 ± 1.0 | 5.8 ± 2.5 | * | 14.9 ± 5.4 | 22.9 ± 8.4 |
|  | 3 | ns | -2.1 ± 2.5 | 0.2 ± 5.0 | ns | 4.9 ± 1.4 | 7.3 ± 4.2 | * | 18.2 ± 5.0 | 28.4 ± 13.6 |
| Trunk LB | 1 | ns | 6.5 ± 3.5 | 8.6 ± 5.4 | ns | 8.7 ± 2.8 | 10.2 ± 4.2 | ns | 21.7 ± 5.5 | 19.9 ± 8.0 |
|  | 2 | ns | 0.2 ± 2.5 | 1.7 ± 2.4 | * | 5.6 ± 2.0 | 8.2 ± 2.6 | ns | 12.8 ± 3.6 | 15.0 ± 3.7 |
|  | 3 | ns | 7.2 ± 3.5 | 11.6 ± 6.0 | * | 11.8 ± 2.8 | 17.3 ± 6.6 | ns | 21.3 ± 3.9 | 24.2 ± 6.6 |
| Trunk AR | 1 | ns | 6.0 ± 3.9 | 5.0 ± 4.6 | ns | 17.8 ± 2.4 | 14.9 ± 4.9 | ns | 42.6 ± 6.6 | 37.0 ± 11.1 |
|  | 2 | ns | 13.7 ± 3.9 | 12.4 ± 5.5 | ns | 15.1 ± 3.0 | 13.8 ± 4.6 | ns | 33.4 ± 8.3 | 36.8 ± 13.6 |
|  | 3 | ns | 13.3 ± 3.8 | 12.9 ± 5.8 | ns | 25.5 ± 3.0 | 24.1 ± 8.2 | ns | 58.6 ± 10.8 | 52.3 ± 17.1 |
| Sho FE | 1 | ns | 51.3 ± 10.6 | 49.3 ± 6.5 | ns | 69.3 ± 7.6 | 61.4 ± 8.5 | * | 192.3 ± 39.4 | 143.8 ± 44.1 |
|  | 2 | ns | 64.9 ± 11.4 | 64.7 ± 8.7 | ns | 72.1 ± 9.7 | 67.1 ± 9.9 | * | 200.8 ± 40.9 | 154.1 ± 45.0 |
|  | 3 | ns | 66.8 ± 11.2 | 67.0 ± 8.8 | ns | 86.0 ± 9.9 | 81.7 ± 10.1 | ns | 233.0 ± 40.4 | 192.8 ± 54.2 |
| Sho AA | 1 | ns | -5.8 ± 5.1 | -6.1 ± 6.8 | ns | 19.3 ± 6.5 | 20.1 ± 5.2 | ns | 76.6 ± 23.6 | 65.6 ± 27.0 |
|  | 2 | ns | 1.4 ± 7.2 | 3.1 ± 9.4 | ns | 25.6 ± 8.8 | 25.6 ± 8.0 | ns | 81.5 ± 30.7 | 69.7 ± 21.3 |
|  | 3 | ns | 3.5 ± 6.9 | 4.0 ± 8.6 | ns | 28.9 ± 9.1 | 32.0 ± 10.5 | ns | 101.7 ± 27.6 | 90.0 ± 24.3 |
| Sho IER | 1 | ns | 22.8 ± 10.0 | 16.4 ± 8.4 | ns | 44.0 ± 7.9 | 41.5 ± 9.2 | * | 151.1 ± 32.3 | 112.5 ± 40.8 |
|  | 2 | ns | 32.6 ± 10.4 | 27.3 ± 9.6 | ns | 32.6 ± 6.7 | 27.8 ± 6.9 | * | 123.3 ± 23.1 | 89.4 ± 34.4 |
|  | 3 | ns | 34.9 ± 9.6 | 29.7 ± 9.7 | ns | 54.2 ± 6.8 | 55.7 ± 10.2 | ns | 180.4 ± 33.8 | 148.9 ± 44.4 |
| Elbow FE | 1 | ns | 92.1 ± 11.9 | 85.4 ± 11.5 | ns | 76.4 ± 10.6 | 73.1 ± 10.2 | * | 274.2 ± 53.8 | 218.5 ± 62.1 |
|  | 2 | ns | 103.6 ± 12.8 | 98.6 ± 12.7 | ns | 81.2 ± 9.6 | 78.8 ± 9.6 | ns | 268.1 ± 47.5 | 226.1 ± 51.4 |
|  | 3 | ns | 103.8 ± 13.2 | 102.3 ± 12.2 | ns | 88.4 ± 11.6 | 87.4 ± 11.3 | ns | 270.3 ± 48.6 | 226.8 ± 55.2 |
| Frm PS | 1 | ns | 40.1 ± 22.5 | 33.8 ± 20.2 | ns | 77.0 ± 15.9 | 78.9 ± 19.0 | * | 308.6 ± 70.4 | 244.7 ± 72.5 |
|  | 2 | ns | 51.3 ± 22.3 | 44.7 ± 20.2 | ns | 51.4 ± 18.2 | 47.1 ± 12.4 | ns | 176.4 ± 57.6 | 149.2 ± 51.7 |
|  | 3 | ns | 51.4 ± 21.7 | 42.7 ± 19.9 | ns | 90.9 ± 17.3 | 85.3 ± 16.4 | ns | 181.8 ± 47.9 | 169.5 ± 62.2 |
| Wrist FE | 1 | * | -18.6 ± 12.4 | -29.1 ± 8.7 | ns | 28.6 ± 6.1 | 31.0 ± 8.4 | * | 136.8 ± 30.4 | 109.3 ± 27.2 |
|  | 2 | * | -11.8 ± 13.8 | -23.5 ± 12.2 | ns | 25.5 ± 8.9 | 32.0 ± 10.3 | ns | 122.3 ± 36.4 | 119.6 ± 37.0 |
|  | 3 | ns | -12.6 ± 11.4 | -22.5 ± 15.3 | ns | 32.3 ± 8.0 | 36.4 ± 14.7 | ns | 123.9 ± 38.6 | 123.0 ± 42.6 |
| Wrist URD | 1 | * | 14.6 ± 7.8 | *23.1 ± 7.1 | ns | 30.9 ± 5.6 | 25.7 ± 7.4 | * | 108.9 ± 39.3 | 77.7 ± 30.1 |
|  | 2 | * | 18.8 ± 7.8 | *26.4 ± 6.8 | ns | 24.7 ± 7.3 | 22.4 ± 7.6 | * | 95.6 ± 23.0 | 69.1 ± 24.1 |
|  | 3 | * | 16.3 ± 7.3 | *24.6 ± 7.0 | ns | 29.7 ± 4.7 | 26.4 ± 5.8 | * | 117.5 ± 28.0 | 88.8 ± 30.8 |

Angular kinematic values include peak angle (degrees), range of motion (degrees), and peak angular velocity (degrees/s) of each movement (M) for trunk flexion/extension (FE), lateral bending (LB), and axial rotation (AR); shoulder (Sho) flexion/extension, abduction/adduction (AA), and internal/external rotation (IER); elbow flexion/extension and forearm pronation/supination (Frm PS); and wrist flexion/extension and radial/ulnar deviation (RUD). For the results of the pairwise comparisons (in column *p*), \* indicates a *p* value less than 0.05, \*\* indicates a *p* value less than 0.005, and ns indicates a *p* value that is not significant. Highlighted table cells also indicate significant differences (red = higher and blue = lower repeated study value).

**Table 5:** Cup Transfer Task angular joint kinematic values, with significant results of the RMANOVAs and pairwise comparisons.

|  | M | Peak Angle (degrees) |  |  | Range of Motion (degrees) |  |  | Peak Angular Velocity (degrees/s) |  |  |
| --- | --- | --- | --- | --- | --- | --- | --- | --- | --- | --- |
|  |  | *p* | Original | Repeated | *p* | Original | Repeated | *p* | Original | Repeated |
| Trunk FE | 1 | ns | -4.4 ± 2.5 | -1.2 ± 6.2 | * | 3.0 ± 1.5 | 4.7 ± 1.9 | ** | 10.7 ± 3.4 | 16.0 ± 4.7 |
|  | 2 | ns | -6.3 ± 2.5 | -2.9 ± 5.7 | ns | 9.1 ± 3.3 | 10.2 ± 2.5 | ns | 23.1 ± 6.8 | 25.2 ± 6.4 |
|  | 3 | ns | -5.6 ± 2.8 | -2.1 ± 6.1 | ns | 9.6 ± 3.1 | 11.2 ± 3.2 | ns | 27.2 ± 7.8 | 31.8 ± 12.2 |
|  | 4 | ns | -5.7 ± 2.7 | -3.0 ± 6.5 | ns | 4.7 ± 2.5 | 6.0 ± 1.5 | ns | 13.0 ± 4.1 | 16.6 ± 5.3 |
| Trunk LB | 1 | ns | -0.4 ± 1.8 | 1.9 ± 3.9 | ** | 4.8 ± 1.9 | 7.7 ± 2.3 | ns | 9.9 ± 3.6 | 12.0 ± 3.4 |
|  | 2 | ns | 0.3 ± 4.1 | 1.9 ± 4.6 | ns | 7.2 ± 2.4 | 9.5 ± 3.6 | ns | 16.5 ± 6.0 | 19.0 ± 8.1 |
|  | 3 | ns | -0.6 ± 3.2 | 2.2 ± 4.1 | ** | 6.2 ± 1.9 | 9.4 ± 3.3 | ns | 15.1 ± 3.7 | 20.9 ± 10.3 |
|  | 4 | ns | -1.1 ± 3.0 | 0.8 ± 4.4 | * | 4.0 ± 1.4 | 5.7 ± 2.0 | ns | 10.8 ± 3.4 | 12.9 ± 5.3 |
| Trunk AR | 1 | ns | 8.9 ± 3.7 | 7.7 ± 5.1 | ns | 9.3 ± 2.5 | 9.2 ± 2.7 | ns | 20.6 ± 4.0 | 21.5 ± 5.2 |
|  | 2 | ns | 17.1 ± 5.0 | 15.7 ± 4.7 | ns | 10.7 ± 2.8 | 11.3 ± 3.4 | ns | 28.1 ± 7.4 | 30.1 ± 9.3 |
|  | 3 | ns | 17.2 ± 5.0 | 16.4 ± 5.0 | ns | 16.7 ± 4.2 | 16.9 ± 4.7 | ns | 39.1 ± 9.8 | 44.2 ± 15.9 |
|  | 4 | ns | 10.3 ± 3.9 | 8.6 ± 4.8 | ns | 7.9 ± 2.4 | 7.2 ± 2.3 | ns | 22.6 ± 7.3 | 22.2 ± 6.7 |
| Sho FE | 1 | ns | 49.2 ± 14.6 | 43.9 ± 9.1 | * | 62.7 ± 13.5 | 50.8 ± 11.4 | * | 142.1 ± 42.8 | 104.5 ± 31.7 |
|  | 2 | ns | 56.8 ± 10.4 | 55.7 ± 7.3 | ns | 30.9 ± 6.2 | 29.6 ± 4.9 | ns | 103.9 ± 29.7 | 77.9 ± 31.7 |
|  | 3 | ns | 57.5 ± 11.0 | 56.4 ± 7.5 | ns | 73.6 ± 10.4 | 66.6 ± 8.9 | * | 228.2 ± 58.8 | 174.3 ± 61.6 |
|  | 4 | ns | 49.5 ± 14.8 | 43.7 ± 10.7 | ns | 29.6 ± 9.0 | 26.4 ± 8.6 | ns | 104.7 ± 25.0 | 95.2 ± 34.5 |
| Sho AA | 1 | ns | -8.1 ± 4.7 | -6.3 ± 6.2 | ns | 27.5 ± 7.1 | 23.7 ± 5.4 | ns | 80.4 ± 23.4 | 65.8 ± 19.7 |
|  | 2 | ns | -1.4 ± 5.9 | -3.4 ± 8.2 | ns | 18.7 ± 5.6 | 16.2 ± 4.2 | * | 63.7 ± 17.1 | 49.3 ± 14.3 |
|  | 3 | ns | 0.1 ± 5.5 | -1.1 ± 7.7 | ns | 28.7 ± 8.6 | 23.9 ± 6.3 | ns | 98.1 ± 34.9 | 79.3 ± 33.0 |
|  | 4 | ns | -13.9 ± 6.9 | -14.4 ± 7.1 | ns | 26.1 ± 6.0 | 21.7 ± 5.8 | * | 74.9 ± 19.4 | 59.3 ± 16.5 |
| Sho IER | 1 | ns | 44.9 ± 14.9 | 35.5 ± 10.9 | ns | 51.5 ± 13.9 | 41.2 ± 10.6 | * | 116.0 ± 57.8 | 74.1 ± 23.8 |
|  | 2 | ns | 43.6 ± 13.8 | 34.8 ± 10.2 | ns | 33.1 ± 7.5 | 28.2 ± 6.5 | ** | 180.2 ± 36.3 | 120.5 ± 43.1 |
|  | 3 | ns | 41.8 ± 13.8 | 32.4 ± 10.1 | * | 49.9 ± 12.2 | 38.8 ± 10.4 | * | 188.8 ± 56.6 | 131.8 ± 49.3 |
|  | 4 | ns | 46.6 ± 14.7 | 37.9 ± 11.8 | ns | 39.5 ± 9.6 | 36.6 ± 8.5 | * | 160.1 ± 39.0 | 120.4 ± 41.4 |
| Elbow FE | 1 | ns | 84.7 ± 12.3 | 78.5 ± 11.7 | ns | 44.6 ± 9.4 | 48.8 ± 10.7 | ns | 173.4 ± 44.5 | 150.6 ± 39.0 |
|  | 2 | ns | 70.6 ± 11.6 | 66.6 ± 11.7 | ns | 60.4 ± 8.1 | 58.8 ± 8.2 | ns | 196.8 ± 30.6 | 174.5 ± 39.5 |
|  | 3 | ns | 93.3 ± 12.9 | 84.0 ± 11.2 | ns | 84.6 ± 9.3 | 78.7 ± 11.8 | * | 281.1 ± 59.3 | 226.6 ± 63.3 |
|  | 4 | ns | 84.7 ± 13.4 | 84.3 ± 11.6 | ns | 48.3 ± 6.0 | 53.5 ± 6.9 | ns | 227.7 ± 43.2 | 213.9 ± 43.6 |
| Frm PS | 1 | ns | 50.7 ± 21.5 | 50.2 ± 17.7 | ns | 31.0 ± 11.5 | 36.3 ± 11.2 | ns | 113.9 ± 24.3 | 125.4 ± 37.1 |
|  | 2 | ns | 36.7 ± 19.5 | 43.2 ± 18.6 | ** | 46.9 ± 12.6 | 62.9 ± 15.0 | ns | 182.4 ± 44.5 | 190.5 ± 69.2 |
|  | 3 | ns | 49.7 ± 22.2 | 38.8 ± 19.9 | ns | 64.2 ± 11.5 | 62.5 ± 17.9 | ns | 196.2 ± 49.1 | 154.7 ± 52.7 |
|  | 4 | ns | 43.3 ± 21.0 | 45.1 ± 19.8 | ns | 46.6 ± 9.7 | 56.6 ± 15.0 | ns | 188.6 ± 67.7 | 184.0 ± 50.4 |
| Wrist FE | 1 | ** | 35.6 ± 11.4 | 22.8 ± 10.5 | ns | 74.2 ± 14.4 | 81.0 ± 16.4 | ns | 283.1 ± 74.0 | 259.6 ± 68.2 |
|  | 2 | * | 28.4 ± 13.6 | 14.8 ± 10.2 | ns | 57.2 ± 7.4 | 55.2 ± 11.5 | ns | 276.5 ± 78.2 | 219.9 ± 87.4 |
|  | 3 | * | 0.9 ± 14.9 | -12.7 ± 10.5 | ns | 34.6 ± 10.9 | 41.9 ± 11.1 | ns | 162.9 ± 65.2 | 138.2 ± 37.0 |
|  | 4 | ** | 44.5 ± 13.6 | 28.6 ± 10.9 | ns | 61.7 ± 10.1 | 58.6 ± 13.1 | * | 299.9 ± 63.0 | 237.5 ± 65.5 |
| Wrist URD | 1 | ns | 24.6 ± 11.4 | 24.3 ± 7.5 | ** | 37.7 ± 8.5 | 26.4 ± 8.2 | ** | 134.9 ± 34.7 | 81.9 ± 26.4 |
|  | 2 | ns | 23.6 ± 9.6 | 24.2 ± 7.6 | ns | 27.7 ± 6.1 | 23.1 ± 7.7 | ** | 122.5 ± 35.3 | 84.1 ± 26.9 |
|  | 3 | ns | 15.8 ± 7.4 | 18.1 ± 8.2 | ns | 25.1 ± 6.2 | 20.6 ± 6.3 | ** | 115.0 ± 35.4 | 73.7 ± 22.1 |
|  | 4 | ns | 26.9 ± 11.7 | 26.5 ± 7.8 | ns | 23.5 ± 6.0 | 20.5 ± 8.8 | * | 126.4 ± 33.9 | 91.8 ± 34.6 |

Angular kinematic values include peak angle (degrees), range of motion (degrees), and peak angular velocity (degrees/s) of each movement (M) for trunk flexion/extension (FE), lateral bending (LB), and axial rotation (AR); shoulder (Sho) flexion/extension, abduction/adduction (AA), and internal/external rotation (IER); elbow flexion/extension and forearm pronation/supination (Frm PS); and wrist flexion/extension and radial/ulnar deviation (RUD). For the results of the pairwise comparisons (in column *p*), \* indicates a *p* value less than 0.05, \*\*
indicates a *p* value less than 0.005, and ns indicates a *p* value that is not significant. Highlighted table cells also indicate significant differences (red = higher and blue = lower repeated study value).

The original and repeated study participants also had similar ROM values in Pasta, although significant differences were found for the Movement 2 trunk flexion/extension ROM and the Movement 2 & 3 trunk lateral bending ROM. However, these differences were quite small (with the largest being 5.3°). In Cups, differences in ROMs were significant in more movements and degrees of freedom (DOFs), as indicated by the shading in **Error! Reference source not found.**. However, the significant trunk ROM differences were quite small (both less than 2°), and the significant shoulder ROM differences were less than the respective original study standard deviations for those DOFs.

The repeated study participants exhibited differences in peak angular velocities in most DOFs in both tasks. The peak angular velocities in the trunk DOFs of repeated study participants were usually greater than those of original study participants, with significant trunk flexion/extension differences in Movement 1 and 2 of Pasta and Movement 1 of Cups. The peak angular velocities in the remaining DOFs of the repeated study participants were usually smaller than for the original study participants, with most significantly lower.

### Eye Gaze

The repeated and original study participants exhibited similar eye fixations, with no significant differences identified in either task, as shown in S5 Table (Pasta) and S6 Table (Cups). Significant eye arrival latency differences were identified in all Grasp phase transitions and the Movement 3 Release phase transition of Pasta, as well as the Movement 3 phase transitions of Cups. No significant eye leaving latency differences were identified in Pasta, but significant differences were identified in the Movement 3 Release transition in Cups.

## Discussion

Measures that were consistent between the original and repeated studies included all hand velocity, grip aperture, and eye fixation results, along with most peak joint angle and ROM results. Although participants in the repeated study took more time to complete each functional task (greater overall duration), similar relative phase durations between the participant groups indicated that the repeated study participants did not spend a disproportionate amount of time in any one phase.

Participants in the original study may have displayed faster performance due to the prior functional task trials that they completed (that is, during task trials where only motion capture or eye tracking data were captured in the original study). This presumption is likely, given that practice has been shown to decrease functional test completion time [14]. The longer phase durations exhibited by the repeated study participants led to both increased eye arrival latencies and decreased eye leaving latencies. Furthermore, their longer movement times resulted in decreased joint angular velocities in shoulder, elbow, forearm, and wrist DOFs.

Learning effects may have also contributed to discrepancies in hand movement measures between the original and repeated study participants. The repeated study participants exhibited an increased number of movement units and increased hand trajectory variability, both of which were likely due to the influence of fewer practice opportunities [15], [16]. Furthermore, increased hand trajectory variability presumably contributed to the repeated study participants’ increased average hand distance travelled. Hand trajectory variances would be expected to be away from, or in avoidance of, obstacles present in all task movements (box walls and the partition in the Cup Transfer Task, and the shelf frames in the Pasta Box Task). Future studies that employ GaMA should standardize the amount of functional task practice opportunities that participants receive.

Task demonstration variations by raters may also have contributed to task duration differences between the two participant groups. Although the same script was used to explain the tasks to participants in each study, small variances in task demonstration speed may have been introduced by the raters. Since the timing of demonstrations is known to influence the resulting pace of participants’ movements [17], a slower demonstration may have contributed to the repeated study’s increase in task duration time. It is recommended that a standard task demonstration video be created and shown to all participants to reduce the possible effects of rater demonstration variation.

The angular kinematic measures revealed offsets in the wrist flexion/extension and ulnar/radial deviation measures of the repeated study participants, likely due to differences in the kinematic calibration pose across the two studies. Such calibration errors are known to be the main limitation of the *Clusters Only* model [13]. In addition, a large standard deviation in trunk flexion/extension was observed for repeated study participants, also likely attributable to errors in the kinematic calibration. That is, the calibration of this DOF depends on how each participant chooses to ‘stand upright’. To limit such deviations in joint angles, the rater must ensure that the participant does not have a bent wrist and is standing as upright as possible, when a kinematic calibration pose is captured.

Further angular kinematics variations were observed between the two participant groups, in both the forearm pronation/supination and wrist radial/ulnar deviation ROMs. Such deviations were introduced by the *Clusters Only* model, which calculates wrist and forearm angles in a manner that is different from other DOFs. This alternative calculation method was chosen because, during the required calibration pose, participants struggled to align their wrist axes of rotation with the global coordinate system, either due to their elbow carrying angle or their inability to supinate their forearm the required amount. As such, the model uses the local coordinate system of the forearm plate to calculate wrist and forearm joint angles. Small misplacements of the forearm marker plate, however, can introduce wrist and forearm joint angle calculation errors. To combat this limitation of the *Clusters Only* model, the rater must take care to align the forearm marker plate with the long axis of the forearm when it is affixed to the participant.

Although little has been done to validate eye tracking and/or motion capture methods in upper limb movement research, many studies have validated motion capture methods for gait measurements [18]. Gait studies commonly revealed that inconsistencies in motion capture marker placement were a large source of anatomical model errors [18]. The *Clusters Only* model used by GaMA attempts to address this issue as it does not require precise individual marker placement, and has been shown to be more reliable than anatomical models [13]; it does, however, introduce its own variability caused by calibration pose inconsistencies. Gait reliability research has also identified intrinsic participant-to-participant variation within a given population and trial-to-trial variation for a given participant [18], [19]. Such variation could, at least partially, also explain movement behaviour differences between the original and repeated data sets of this study.

### Limitations

Given that this study manipulated numerous experimental factors when comparing the visual and movement measures of two groups of non-disabled participants, it had limitations. It was infeasible for this research to determine the degree to which these factors (different participants, sites, equipment, raters, and task experience opportunities) affected movement measure variation. Additional research on the effects of training could shed more light onto whether or not the amount of practice fully explains the difference in results between the two studies. Although assessment of inter-site/inter-rater reliability of GaMA using the same participant group would also provide valuable information by reducing the effects of inter-participant variability, for this study, a new participant group presented an opportunity to analyze a wider range of normative behaviour; an important consideration when designing an assessment tool to be used to characterize functional impairments.

## Conclusions

Overall, the results of the repeated study were similar to those obtained by Valevicius et al. and Lavoie et al. [7]–[9]. Most hand movement, angular joint kinematic, and eye gaze results exhibited by participants in the repeated study were consistent with those observed in the original study. Most significant differences between the results could be explained by the amount of practice that participants in the two studies received, demonstration variations introduced by the rater, and the limitations of the *Clusters Only* kinematics model. Due to its demonstrated reproducibility, it is expected that, in the future, GaMA can serve as a reliable and informative functional assessment tool across different sites and for individuals with sensorimotor impairments in the upper limb.

## Acknowledgements

We thank Quinn Boser, Aida Valevicius and Thomas R. Dawson for assistance with the original data set analysis.

## Supporting Information

**S1 Text.** Detailed statistical analysis.

**S2 Table.** Phase duration values for the Pasta Box Task and Cup Transfer Task, with significant results of the pairwise comparisons. For the results of the pairwise comparisons (in column p), * indicates a significant p value less than 0.05, ** indicates a p value less than 0.005, and “ns” indicates a p value that is not significant.

**S3 Table.** Pasta Box Task hand movement values for each movement segment, with significant results of the pairwise comparisons. For the results of the pairwise comparisons (in column p), * indicates a significant p value less than 0.05, ** indicates a p value less than 0.005, and “ns” indicates a p value that is not significant.

**S4 Table.** Cup Transfer Task hand movement values for each movement segment, with significant results of the pairwise comparisons. For the results of the pairwise comparisons (in column p), * indicates a significant p value less than 0.05, ** indicates a p value less than 0.005, and “ns” indicates a p value that is not significant.

**S5 Table.** Pasta Box Task eye movement values, with significant results of the pairwise comparisons. For the results of the pairwise comparisons (in column p), * indicates a significant p value less than 0.05 and “ns” indicates a p value that is not significant.

**S6 Table.** Cup Transfer Task eye movement values, with significant results of the pairwise comparisons. For the results of the pairwise comparisons (in column p), * indicates a significant p value less than 0.05 and “ns” indicates a p value that is not significant.

